# Variational Log-Gaussian Point-Process Methods for Grid Cells

**DOI:** 10.1101/2023.03.18.533177

**Authors:** Michael Everett Rule, Prannoy Chaudhuri-Vayalambrone, Marino Krstulovic, Marius Bauza, Julija Krupic, Timothy O’Leary

## Abstract

We present practical solutions to applying Gaussian-process methods to calculate spatial statistics for grid cells in large environments. Gaussian processes are a data efficient approach to inferring neural tuning as a function of time, space, and other variables. We discuss how to design appropriate kernels for grid cells, and show that a variational Bayesian approach to log-Gaussian Poisson models can be calculated quickly. This class of models has closed-form expressions for the evidence lower-bound, and can be estimated rapidly for certain parameterizations of the posterior covariance. We provide an implementation that operates in a low-rank spatial frequency subspace for further acceleration, and demonstrate these methods on experimental data.

## Introduction

Grid cells in the hippocampal formation modulate their firing rates as a periodic function of location (Hafting et al., 2005; Rowland et al., 2016). Some grid cells are also modulated by head direction (Sargolini et al., 2006; “conjunctive cells”), and recent studies have found mode subtle dependence on head direction and (Gerlei et al., 2020) and landmarks (Keinath et al., 2018; Krupic et al., 2018) even in non-conjunctive cells. Exploring these relationships requires efficient statistical estimators to compare changes in the spatial dependence of grid-cell activity across conditions.

Standard approaches to spatial statistics have limitations. Grid-cell firing-rate maps are often estimated using a Gaussian kernel-density smoother (e.g. Hafting et al., 2005; Langston et al., 2010; Brandon et al., 2011; Killian et al., 2012). Naïve smoothing approaches remain noisy when data are limited, do not provide a quantification of uncertainty, cannot adapt to inhomogeneous spatial sampling, and cannot take advantage of the periodic structure of grid-cell firing. Conversely, approaches based on spatial autocorrelations (e.g. Hafting et al., 2005, many others) reduce noise by averaging over space, but cannot be applied to single grid fields. Gaussian-Process (GP) estimators are a promising solution to these challenges. They offer a principled, Bayesian approach to estimating firingrate maps. They incorporate assumptions to improve statistical efficiency, and provide a posterior distribution that quantifies uncertainty.

However, open challenges remain in applying existing algorithms to exploratory analysis of large grid-cell datasets. Bayesian priors suitable for grid cells have not been described in the literature, and existing implementations are either limited to specific kernels or are too computationally intensive for large datasets. We resolve both of these issues, and illustrate practical benefits of Gaussian-process methods compared to non-Bayesian estimators.

We briefly review Gaussian process methods in neuroscience, then present (1) a tutorial on applying Gaussian processes to grid-cell data; (2) a technical review of approximate inference algorithms; (3) applications of these methods on example data.

## Background

Gaussian processes generalize the multivariate normal distribution to a distribution over functions (MacKay, 1998; Rasmussen, 2003; Keeley and Pillow, 2018). They are a natural candidate for describing neuronal tuning as a function of continuous variables, and have emerged as the gold-standard for analyzing neuronal activity in the low-data regime. Many algorithms have been developed for capturing the relationship between neural activity and other variables, or for inferring latent neural states (Yu et al., 2009; Rad and Paninski, 2010; Park et al., 2014; Frigola et al., 2014; Wu et al., 2017; Zhao and Park, 2017; Duncker and Sahani, 2018; Brandman et al., 2018; Rule et al., 2019; Keeley et al., 2020; Jensen et al., 2020, 2021).

Formally, a Gaussian-process distribution is specified by its mean function *μ* (*x)* and two-point covariance function Σ (***x, x***^′^), which are analogous to the mean vector ***μ*** and covariance matrix **Σ** of the multivariate normal distribution (see MacKay, 1998; Rasmussen, 2003; Keeley and Pillow, 2018 for a thorough introduction). In computation, however, Gaussian processes are almost always represented in terms of a finite-dimensional approximation. We will use the finite-dimensional notation **z** ∼ 𝒩 (***μ*, Σ**), with the understanding that this represents a particular finite-dimensional projection of our Gaussian process model.

Previous works have described Gaussian process methods for place and grid cells (e.g. Rad and Paninski, 2010; Wu et al., 2017; Savin and Tkacik, 2016). However, we encountered practical deficiencies when applying these methods to grid cells in large arenas. Computational efficiency is paramount for exploratory analyses of large datasets. While scalable solutions exist, the fastest methods require spatial covariance priors that can be described in terms of nearest-neighbor interactions (Rad and Paninski, 2010; Cseke et al., 2016) or a product of rank-1 separable kernels (Savin and Tkacik, 2016). This is not ideal for grid cells, which can display spatial correlations between response fields separated by several centimeters, and which cannot be decomposed into a product of 1D kernels. Recent works have developed ways to approximate the Gaussian process covariances that support fast calculations, while remaining expressive (Jensen et al., 2021). We elaborate upon these ideas, with a particular focus on grid cells, and introduce some new numerical approaches.

Specifically, the new contributions of this manuscript are (1) Tools for designing GP priors that take advantage of the local spatial topography of grid cells; (2) Efficient and expressive variational Bayesian methods; (3) Numerical algorithms with good performance on consumer-grade hardware; (4) A Python reference implementation and example application to grid-cell data.

## Results

We will first review log-Gaussian Poisson models of neural spiking in the context of inferring a grid-cell firing-rate map. These combine a Gaussian prior on (log) firing rate with a Poisson likelihood for spikes. We review numerical approaches for finding Bayesian posterior, and discuss suitable priors for grid cells, and finally demonstrate applications on example data.

### An example experiment

Throughout this text, we will demonstrate Gaussianprocess methods on data from Krupic et al. (2018), which have also presented in Chaudhuri-Vayalambrone et al. (2023). Figure 1 illustrates a spatial-navigation experiment (Krupic et al., 2018) in which a rat foraged in a 2 m ×1 m environment (Fig. 1a). Spike counts *y*_*t*_ from a grid cell in entorhinal cortex, along with position ***x***_*t*_ = {*x*_1;*t*_, *x*_2;*t*_ }^⊤^, were recorded in 20 ms bins, yielding time series ***X*** = {***x***_1_, .., ***x***_T_ }^⊤^ and ***y*** = {*y*_1_, .., *y*_T_ }^⊤^ with T samples. Throughout this manuscript, we will denote scalars as lower-case letters “*x*”, column vectors as bold lower-case letters “***x***”, and matrices as bold capital letters “***X***”.

**Figure 1:**
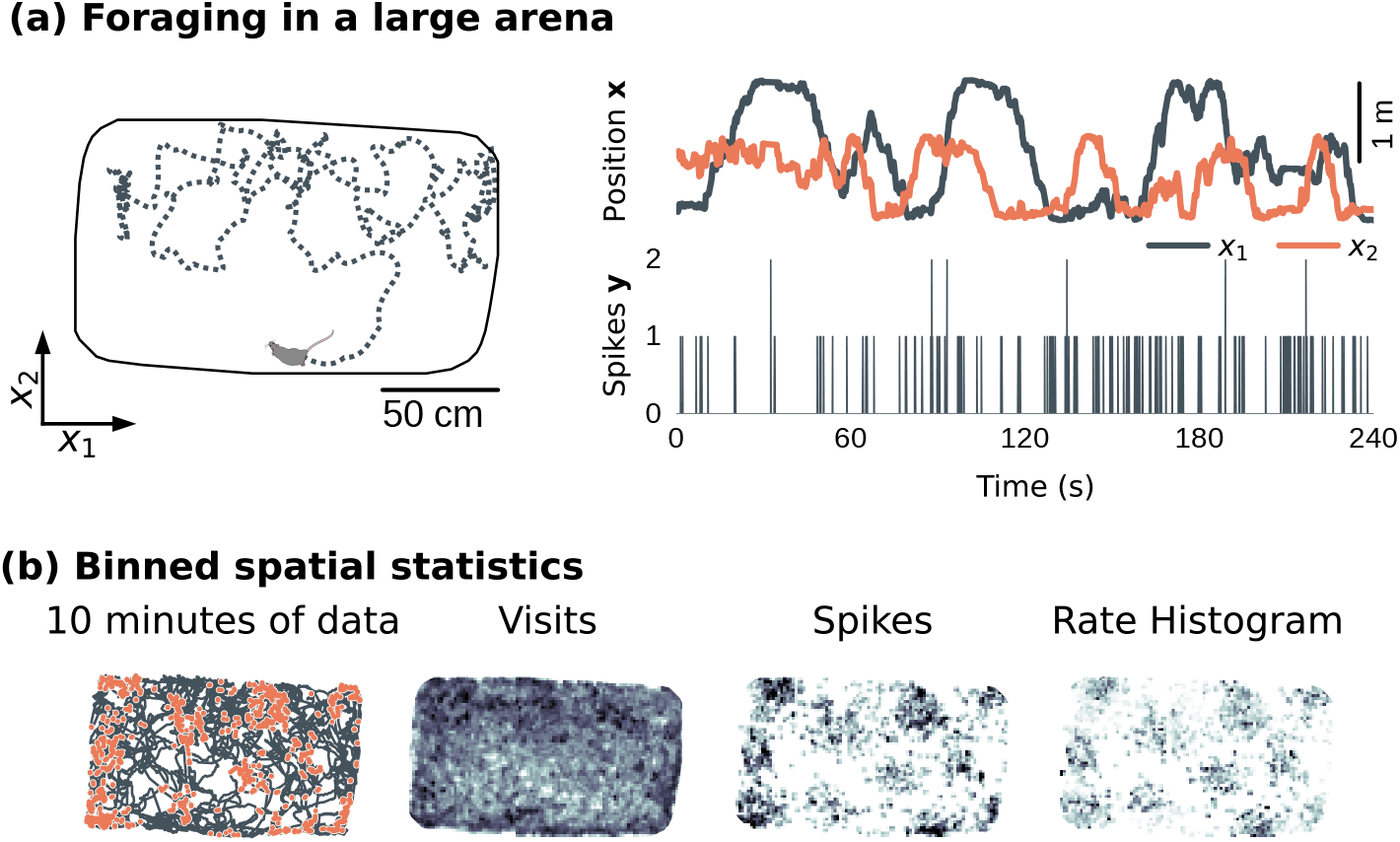
An example experiment. (a) In this experiment, a rat foraged in a 2 m× 1 m open environment (left). The rat’s position over time “***x***” (right, top), as well as spike counts “***y***” from a single neuron in entorhinal cortex (right, bottom) were recorded (data from Krupic et al., 2018). **(b)** A firing-rate histogram (right, ***k*** /***n***) can be estimated by dividing the total number of spikes tallied at each location “***k***” (left) by the number of visits to each location “***n***” (middle). (Color scales are not quantitative.)

The resulting spatial data consists of a map of the number of times the rat visited each location, and the number of spikes observed during each visit. These can be summed on a spatial grid to form occupancy and spike-count histograms, which can be combined to yield a firing-rate histogram (Fig. 1b, 4a). In Figure 1, we binned data on a 88 × 128 grid.

### Estimating a smoothed log-rate map

Our approach will follow variational inference for Gaussian-process generalized linear models as outlined in Challis and Barber (2013). We consider “latent” Gaussian processes, whose values are observed through a firingrate nonlinearity and neuronal spiking activity. We model the log-firing-rate z (***x***) (Fig. 4b) as a Gaussian process, and spiking events as conditionally Poisson (Figure 2a). This model is sometimes called a log-Gaussian Cox process, after David Cox (Cox, 1955). It captures both correlations and over-dispersion in the covariance structure of z(***x***).

**Figure 2:**
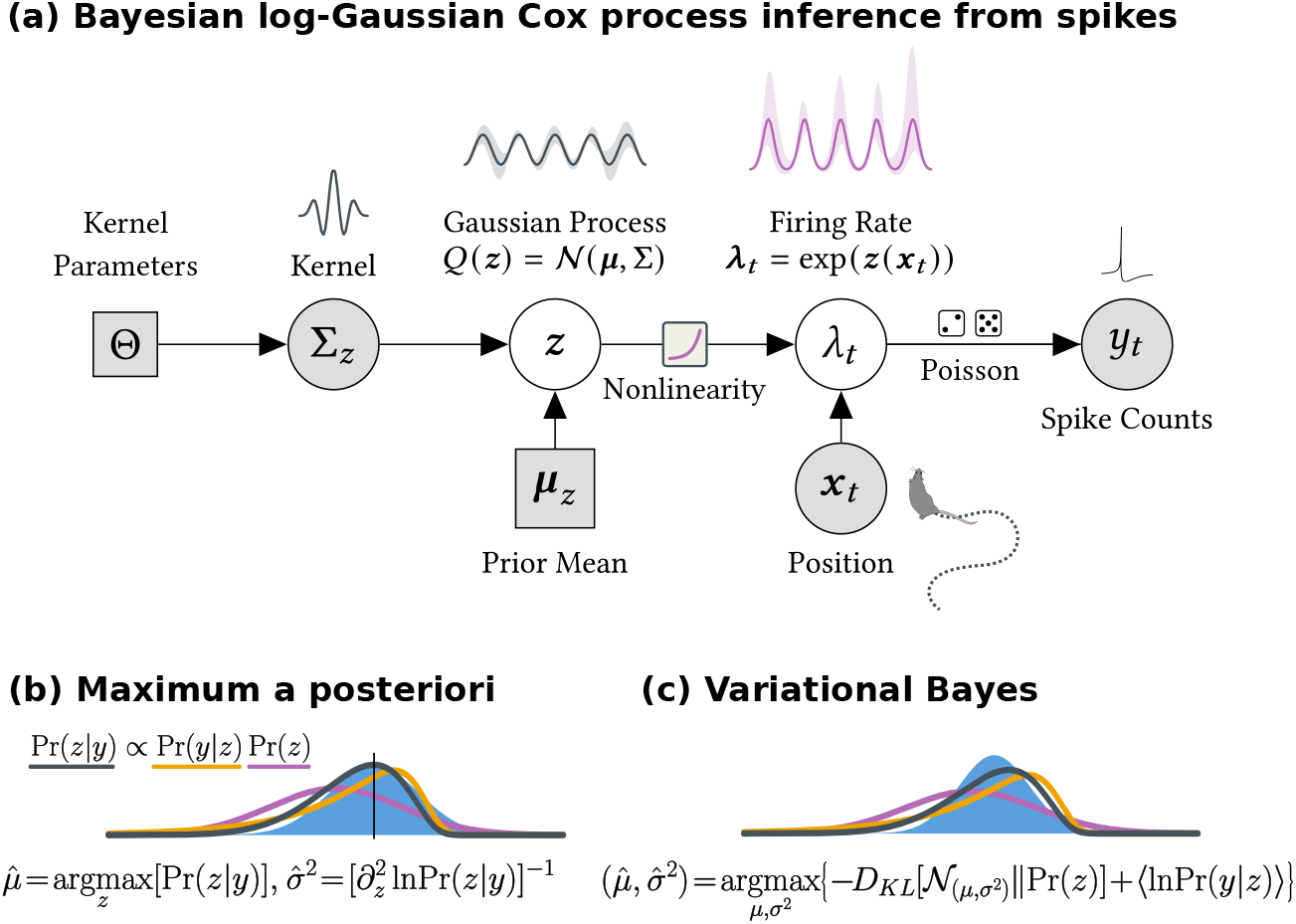
A Bayesian model for firing-rate maps with spiking observations. **(a)** A graphical diagram of the inference procedure. The prior mean and kernel are set externally. A log-Gaussian process parameterizes the inferred firing-rate map. Spiking observations are explained in terms of spatial tuning to location. (b) The posterior distribution over the log-rate map ***z*** is difficult to calculate directly. The maximum *a posteriori* (Methods: *Finding the posterior mode*) estimator approximates Pr (***z***) as Gaussian, with mean equal to the posterior mode, and covariance taken from the curvature at this mode (Methods: *Connection to the Laplace Approximation*). **(c)** Variational Bayesian inference finds a multivariate Gaussian model for the posterior on ***z*** by maximizing a lower-bound on the model likelihood. This can be more accurate when the posterior is skewed, and the same lower bound can be used to select hyperparameters.

We model spike counts within a small time bin Δ*t* as Poisson distributed with firing rate *λ*(***x***) = exp[*z*(*x*)]:

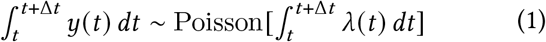

The choice of an exponential firing-rate nonlinearity *λ*= exp (*z*) is useful for obtaining closed-form solutions in variational inference. For simplicity, we will choose time coordinates such that Δ*t* = 1 and omit it going forward. The log-likelihood of observing spike count *y* given rate *λ* is then:

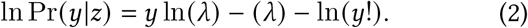

The overall likelihood of all spiking observations ***y*** depends on the log-firing-rate map *z*(***x***), and the animal’s trajectory over time ***X***. We assume that the spiking observations are independent conditioned on the log-rates ***z***, so that the likelihood of the overall dataset Pr (***y***| ***z, X***) factors as

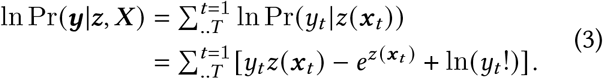

For numerical implementations, we model the function *z* (***x***_*t*_) as a vector ***z*** = {*z*_1_, .., *z*_*M*_ }, where each *z*_*m*_ reflects the value of *z* (***x***_*m*_) at one of *M* spatial locations. To make the notation easier to read in these derivations, we will interpret each location *z*_*m*_ as a piecewise-constant model of the firing-rate map in a small region of the environment with value *z* (***x***) ≈*z*_*m*_ if ***x*** ∈??_*m*_ (in practice we use linearly interpolated binning for improved resolution; Methods: *Binning data*).

We aggregate time points that fall in the same spatial bin, since these share the same log-rate *z*_*m*_ (this is a form of pseudo-point method; Quinonero-Candela and Rasmussen, 2005). We refer to individual bins by a single index *m* ranging from 1 to *M*. We denote the tallies of visits to each bin as ***n*** = {*n*_1_, .., *n*_*m*_}^⊤^ and the tallies of spikes in each bin as ***k*** = {*k*_1_, .., *k*_*m*_}^⊤^:

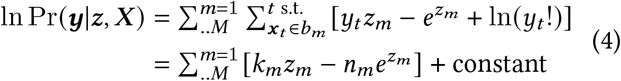

Since ln (*y*!) does not influence the gradient of (4) with respect to **z**, we ignore it when optimizing **z**. Having combined data from repeated visits to the same location, the likelihood in Equation (4) can then be written in vector notation as:

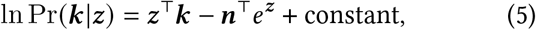

This is the log-likelihood of observations ***k*** given **z**.

This observation model has the same form as a PointProcess Generalized-Linear Model (PP-GLM; e.g. Paninski, 2004; Truccolo et al., 2005; Truccolo, 2016). However, adjusting **z** to maximize (5) alone will lead to overfitting. Instead, one can obtain a smoothed map by taking a Bayesian approach.

We can encode constraints like smoothness or periodicity in our choice of the prior Pr (**z**). We use a multivariate Gaussian prior ***z*** ∼ 𝒩 (***μ***_*z*_, **Σ**_*z*_), which has the logprobability density

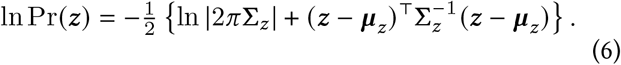

Summing the log-likelihood (5) and log-prior (6) yields an expression for the log-posterior of ***z*** (up to constant terms):

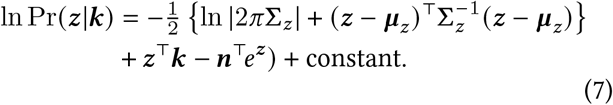

When the dimension of ***z*** is large, estimating (7) via sampling or evaluating it on a grid is infeasible. Instead, we approximate the posterior as a multivariate Gaussian distribution.

### Covariance kernels for grid cells

Throughout this manuscript, we assume that the prior covariance between two points **Σ**_*z*_ (***x***_1_, ***x***_2_) depends only on the displacement between them. In this case, the prior covariance takes the form of a convolution kernel. Since we evaluate our rate map on a rectangular grid, and since the prior covariance is a convolution, **Σ**_*z*_ is a circulant matrix and products like 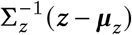 can be computed using the Fast Fourier Transform (FFT) in 𝒪 (*M* log (*M*)) time. (Note: when implementing convolutions via the FFT, it is important to add spatial padding equal or larger than the kernel’s radius, to avoid erroneous correlations from the periodic boundary.)

How should one select **Σ**_*z*_? In Gaussian process regression, the covariance kernel describes how correlated (or anticorrelated) two points ***x***_*i*_ and ***x*** _*j*_ in the inferred rate map are expected to be, as a function of the displacement between them: [**Σ**_z_]_*i j*_ = 𝒦 (***x***_*i*_ −***x*** _*j*_). For any collection of spatial locations, the **Σ**_*z*_ induced by the kernel needs to be a valid covariance matrix; **Σ**_*z*_ must be positive definite: It should be symmetric, real-valued, and have all positive eigenvalues. For our inference procedure to be sensitive to grid cell’s periodicity, our kernel needs a periodic structure. A hexagonal map with period *P* and orientation *θ*_0_ can be defined as the sum of three cosine plane waves, rotated at *π*/3 radians from each-other:

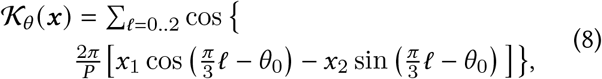

where ***x*** = {*x*_1_, *x*_2_ }∈ℝ^2^. The ideal grid (8) is a valid kernel function: It is symmetric, and its Fourier transform consists of all non-negative real coefficients.

We also use a radially symmetric kernel (Fig. 3a-3, 3a-4) for analyzing grid-cell period in an orientation-agnostic manner. We can construct a radial kernel by considering a ring of spatial-frequency components *ξ* = *ρe*^*iω*^ that match the spatial frequency *ρ* = 1 /*P* of the grid, or, equivalently, a radially-averaged version of (8). In this spatial domain, this kernel is the zeroth-order Bessel function of the first kind,

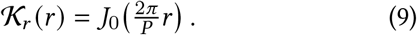

This kernel is more general: It does not require a fixed, global grid orientation, and can be applied to cells with fields separated by a characteristic distance, but no global lattice (as seen in the entorhinal cortex of bats, Ginosar et al., 2021—although in 3D the radial kernel (9) takes the form 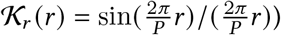.

**Figure 3:**
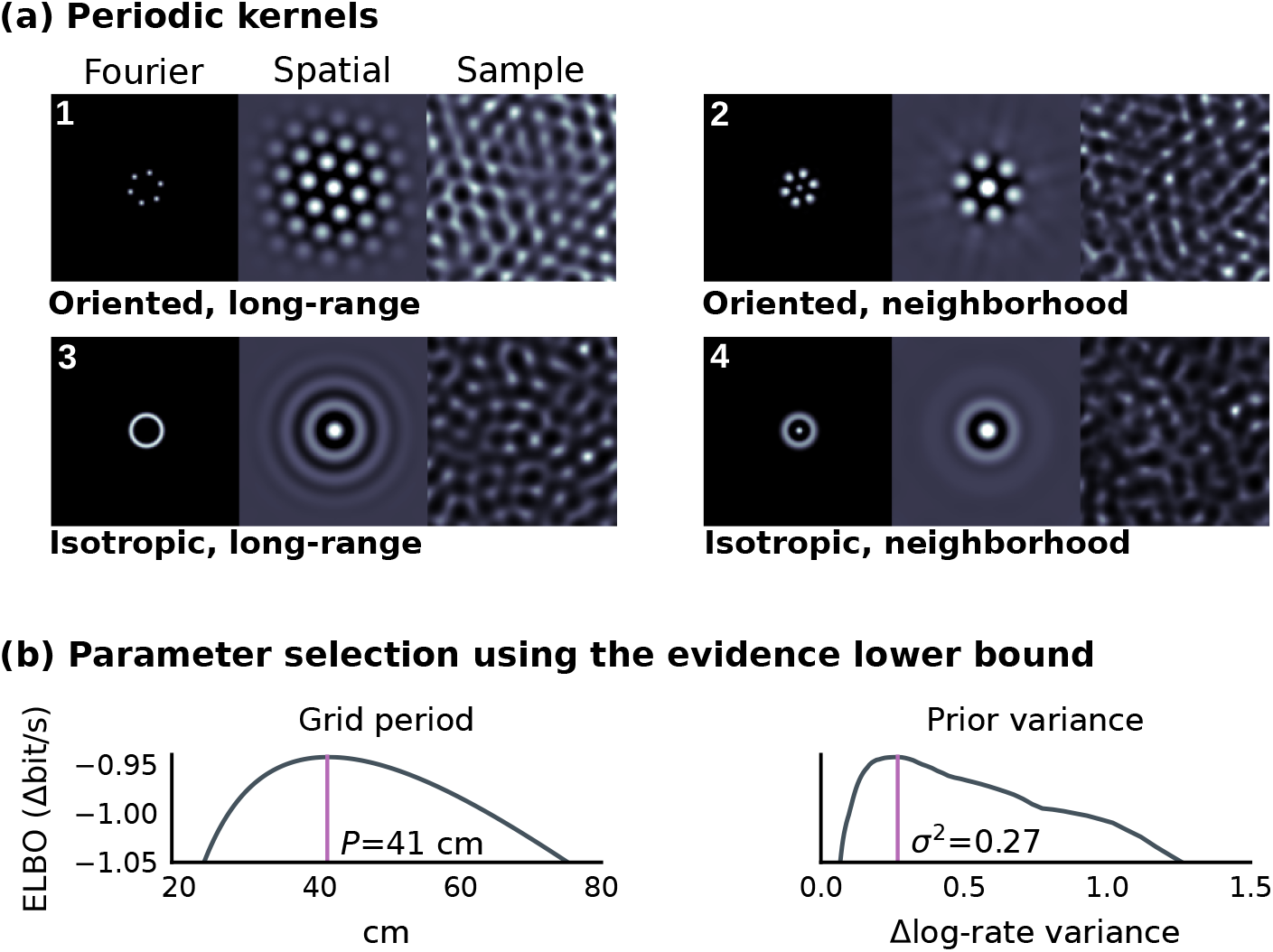
Periodic priors to infer gridcell maps. **(a)** Periodic kernels suitable for grid cells. Each plot shows the kernel’s 2D Fourier spectrum (left), spatial domain representation (center), and an example rate map sampled from the kernel (right). (1,2): Oriented kernels are selective for the grid cell’s preferred spatial orientation. (3,4): Radial kernels based on the Bessel function include no prior assumptions about grid orientation. **(b)** Kernel parameters, like grid cale, can be selected by choosing the kernel that gives the best Evidence Lower Bound (ELBO) after fitting the posterior rate map. Shown here are the loss functions for the period and variance of an oriented grid kernel (Fig. 3a-2) for the cell in Figure 1.

**Figure 4:**
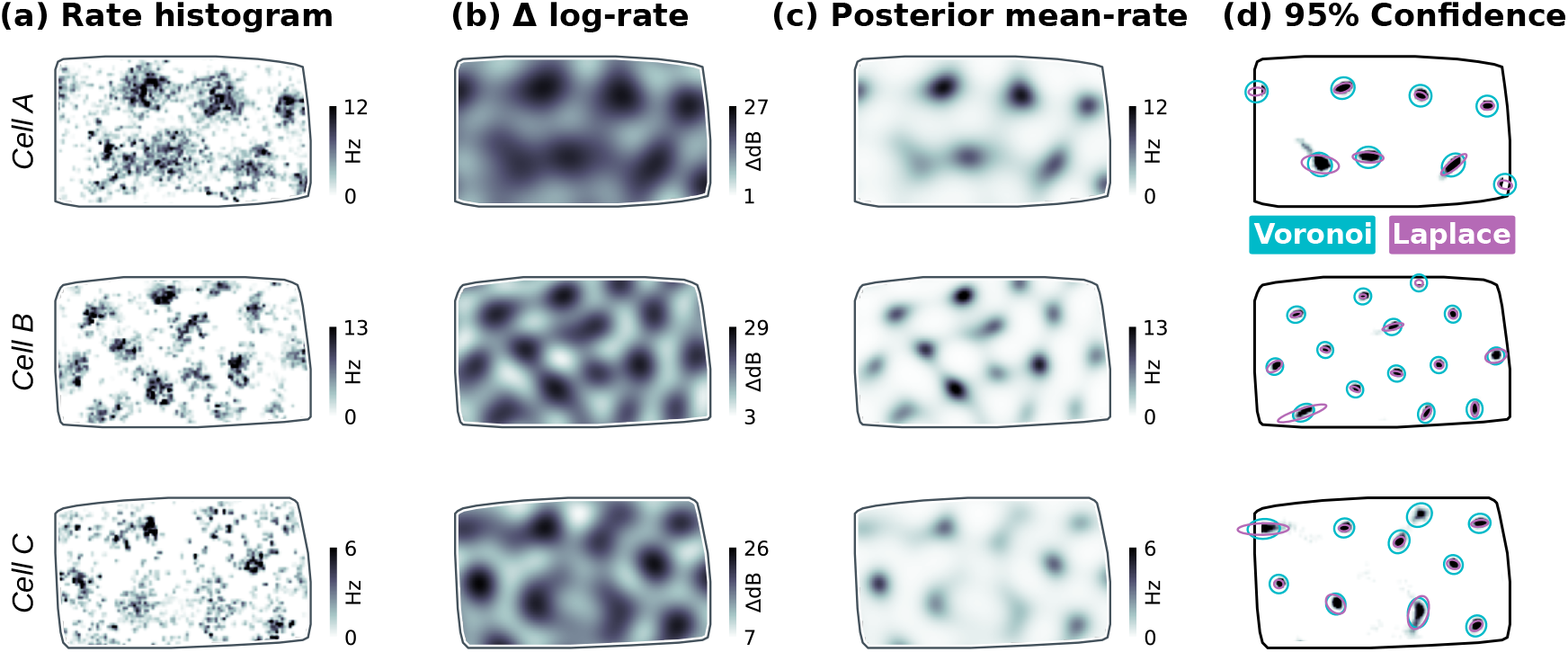
Inferring grid-cell firing-rate maps with LGCP regression. **(a)** Rate histograms from three example cells from Krupic et al. (2018). **(b)** Posterior log-rate map from LGCP inference using the optimized grid-cell kernel (Fig. 3a-2). (Background variations in log firing-rate not included). **(c)** Expected firing rate calculated using (16) from the variational posterior. **(d)** 95% confidence intervals for field location calculated either using a locally-quadratic approximation (purple) (26) or sampling (teal) within each grid-field’s Voronoi region (all points closer to a given field than any other, an no further away than 70% of the grid period), overlaid on the probability density of grid-field peaks (shaded).

The zeros of (9) provide rule-of-thumb cutoff radii for various degrees of spatial interaction: The first zero corresponds to single fields, the second to an inhibitory surround, and the third to nearest-neighbor interactions. In this work, we truncate kernels to nearest-neighbor interactions at *r*_*c*_ = *k*_3_*P* / (2*π*), where *k*_3_ ≈8.65 is the third zero of *J*_0_. We apply a circular window 𝒲 (Δ***x***)= *ϑ* (|Δ***x*** |−*r*_*c*_) (*ϑ* is the Heaviside step function), remove high spatial frequencies from the kernel by applying a 2D Gaussian-blur 𝒦_*σ*_ with radius *σ* = *P* /*π*, and finally truncate any resulting negative Fourier coefficients to zero. This heuristic procedure provided good spatial locality while limiting the kernel to the spatial frequencies of interest; We do not exhaustively compare possible kernels here, but do provide other windowing methods and options to control kernel anisotropy in the reference implementation.

We introduce scale 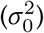 and constant offset (*c*) parameters to control the kernel’s marginal variance, and the variance assigned to the mean-log-rate component, respectively. Using either a grid or radial kernel as a base kernel (𝒦_0_) we define the parameterized kernel 𝒦_**Θ**_ as:

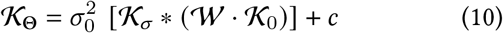

where ∗ denotes convolution and · pointwise multiplication. We discuss hyperparameter selection in Section *Optimizing kernel hyperparameters*.

Generally, one can construct suitable kernels by computing the autocorrelation of a prototype firing-rate map, averaging to achieve any desired symmetries, and applying desired spatial or spectral windowing. If the kernel is defined as a convolution over a regular grid, these operations can be computed quickly using the FFT. Since any product, convolution, or non-negative linear combination of positive-definite kernels is also positive definite, complicated kernels can be constructed out of simple primitives.

### Variational Inference

One can optimize the log-posterior (7) in ***z*** to obtain a smoothed firing-rate map. This is known as the Maximum *a posteriori* (MAP) estimator (Fig. 2b; Methods: *Finding the posterior mode*). The MAP estimator allows us to specify prior assumptions (e.g. smoothness and periodicity) by selecting the appropriate prior covariance **Σ**_*z*_. However, it is important to assess our confidence in the resulting rate map, and to have a formal way of checking whether our prior is reasonable. Variational Bayesian methods provide a formal way to approximate posterior uncertainty, in the form of a Gaussian-process covariance function.

In variational Bayesian inference (Fig. 2c), we approximate the true posterior with a simpler distribution “*Q*_***ϕ***_ (***z***) “ defined by some parameters ***ϕ***. We use a multivariate Gaussian approximation here, so ***ϕ*** = (***μ*, Σ**) and *Q*_***ϕ***_ has the log-probability density

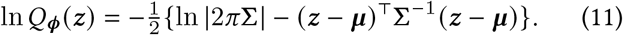

Variational inference selects ***ϕ*** by maximizing a quantity called the evidence lower bound. This is equivalent to simultaneously minimizing the Kullback-Leibler divergence “***D***_KL_” from the prior to the posterior, while maximizing the expected log-likelihood (5) under *Q*_***ϕ***_ :

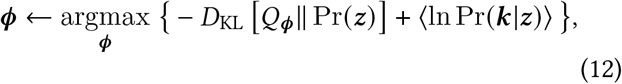

where ⟨·⟩ denotes expectation with respect to *Q*_***ϕ***_ .

The first term in (12) reflects the information gained by revising our estimates of ***z*** compared to our prior beliefs. Since both *Q*_***ϕ***_ (***z***) and Pr (***z***) are multivariate Gaussian, this term has the closed form:

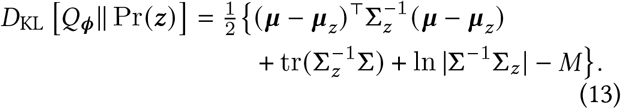

The second term in (12) is the expectation of our Poisson observation model (5) with respect to *Q*_***ϕ***_ :

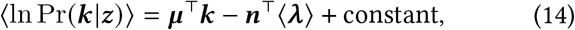

where we abbreviate exp(***z***) as ***λ***. We can write the overall objective “ ℒ” to be maximized as:

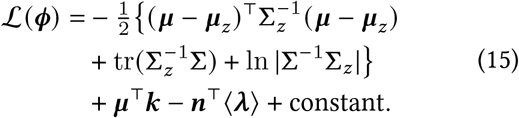

The term “ 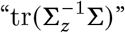 ”; encourages the posterior covariance to be close to the prior, and the term “− ln |**Σ**| “ encourages the posterior to have large entropy.

A convenient property of the log-Gaussian-Poisson model is that the expected firing rate ⟨***λ***⟩ (Fig. 4c) required to calculate (15) has a closed form. Since we have assumed a multivariate Gaussian distribution for ***z***, and since *λ* = exp (***z***), the firing-rate ***λ*** is log-normally distributed. The expectation ⟨***λ***⟩ is the mean of this lognormal distribution, and has the closed-form expression

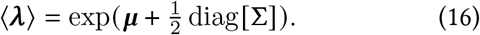

To simplify notation, we define “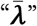” as the expected rate (with dependence on ***μ*** and **Σ** implicit), corrected for the number of visits in each location, i.e. 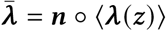. We discuss numerical approaches for calculating (16) briefly in the next section, and in more detail in Methods: *Calculating the expected firing rate*.

### Optimizing the variational posterior

With these preliminaries out of the way, we now consider the derivatives of (15) in terms of ***μ*** and **Σ**. These can be computed using modern automatic differentiation tools (e.g. Jax; Bradbury et al., 2018). However, substantial speedups are possible by considering the analytic forms of the derivatives, and identifying simpler ways to calculate them. The gradient and Hessian of (15) with respect to ***μ*** are

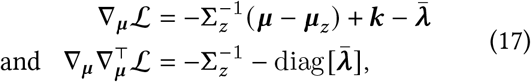

respectively. The derivative of (15) in **Σ** is more involved (Methods: *Derivatives*, Equations (33)-(35)):

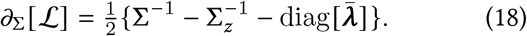

Optimizing the full *M* ×*M* posterior covariance is impractical. Typically, one chooses a simpler parameterization. Combinations of low-rank factorizations and Toeplitz or circulant matrices are common (Jensen et al., 2021). In our case, an *exact* low-dimensional parameterization of the variational posterior covariance is available (Seeger, 1999; Challis and Barber, 2013 Eq. (10)). Note that the stationary point of (18) occurs when 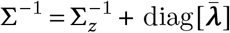. This means that *all* variational posterior covariance matrices can be parameterized by a diagonal update diag 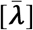 to the prior precision matrix 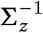. We parameterize this update by the vector ***q*** = {*q*_1_, .., *q*_*M*_ }, and seek a self-consistent solution 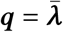:

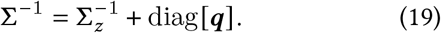

This models the posterior precision as a sum of the prior precision, plus information provided by observations at each location.

We obtain the gradient of ℒ in ***q*** from (18) and (19) using the chain rule (Methods: *Derivatives*, Equations (35)-(37)):

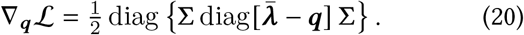

This gradient is zero when 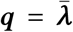. If **Σ** is full rank, this zero is unique, and one may optimize ***q*** by ascending the much simpler gradient 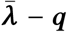, which has the same fixed point.

We maximize the evidence lower bound (15) by alternatively updating ***μ*** and ***q***. Updates to ***μ*** are similar to finding the MAP estimator. We optimize the posterior covariance for fixed ***μ*** via an iterative procedure that amounts to setting 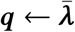 repeatedly (Methods: *Iteratively estimating* ***q***).

There is one remaining difficulty to address. Calculating the expected firing rate (16) requires computing diag [**Σ**]. These are the marginal variances of the firing-rate at each location. For the parameterization in Equation (19), one must compute

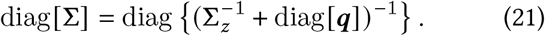

We calculate this using a low-rank approximation of the posterior covariance in Fourier space (Methods: *Working in a low-rank subspace*).

We summarize all steps of this iterative procedure in pseudocode in Algorithm 1. The key takeaways regarding the numeric implementation are this: (1) The posterior mean can be optimized readily using Newton-Raphson iteration, in much the same way as one might estimate the posterior mode for a log-Gaussian-Poisson generalized linear model; (2) The ideal parameterization of the variational posterior covariance takes the form of a diagonal update to the precision matrix, which reflects the amount of information available at each spatial location. This can be updated by a straightforward fixed-point iteration reminiscent of the Laplace approximation (Methods: *Laplace Approximation*).

### Optimizing kernel hyperparameters

The prior covariance kernel in Equation (10) depends on unknown hyperparameters **Θ**: period “*P*”, scale “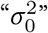”, and mean offset “*c*” (and, for grid kernels, orientation “*θ*_0_”). The variational Bayesian framework provides a principled way to optimize these. To evaluate the quality of hyperparameters, one first optimizes the variational posterior using the kernel determined by **Θ**. At the optimized *Q*_***ϕ***_ (**z**), Equation (12) lower-bounds the likelihood of the data for the chosen hyperparameters. This allows one to compare the quality of different choices of **Θ** (e.g. Fig. 3b). We optimized **Θ** using a hill-climbing grid search, starting from a heuristic guess (*Methods:* Initializing parameters).

### Sampling spatial statistics

Once obtained, the Gaussian-process posterior can be used to sample the distribution of likely firing-rate maps. For example, one may wish to obtain the probability distribution of the peaks of individual grid fields (Fig. 4d).

Given a Gaussian posterior ***z*** ∼**𝒩** (***μ*, Σ**), one can draw samples as ***z*** ←***μ*** +**Σ**^1/2^***η***_*M*_ where ***η***_*M*_ ∼ **𝒩** (0, I_*M*_) is a vector of *M* Gaussian random numbers with unit-variance and zero-mean. However, obtaining **Σ**^1/2^ is impractical for large *M*. Sampling in the low-rank (***D*** *<M*) space 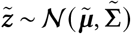 is efficient (Methods: *Working in a low-rank subspace*). Samples can be drawn as

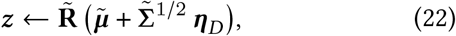

where 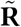 maps samples from the low-rank subspace into the full (spatial) representation, and is described in (41) and (42). The factor 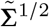 can be calculated as 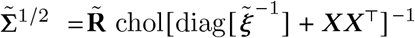, where 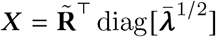 (Methods: *Calculating the expected firing rate*; (43)).

Figure 4d uses sampling to visualize uncertainty in grid-field locations. We generated a peak-density map by plotting the fraction of samples that contain a local maximum within a radius of *P* /2, where *P* is the grid cell’s spatial period. We segmented the arena into Voronoi cells associated with each grid field (out to a maximum radius of 70% P), and calculated 95% confidence ellipses by fitting a 2D Gaussian to each segmented grid field’s peak distribution.

### Peak-location confidence intervals

For well-identified grid fields, one can calculate confidence intervals from the posterior distribution using a locally quadratic approximation. Consider a local maximum in the posterior mean ***μ*** (***x***) at location ***x***_0_. How much does ***x***_0_ change if a perturbation ***ε*** (***x***) ∼ 𝒩 [0, **Σ**(***x, x***^′^)], sampled from the posterior covariance, is added to ***μ*** (***x***)?

This can be calculated via a Taylor expansion in Δ_***x***_ = ***x*** −***x***_0_ of ***μ*** (***x***) at ***x***_0_. The slope at ***x***_0_ is zero, since it is a local maximum, so a Taylor expansion out to second order has only 0^th^and 2^nd^-order (curvature) terms. The curvature in ***x*** is defined by the Hessian matrix 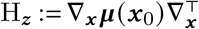. Out to second order our grid-field log-firing-rate is:

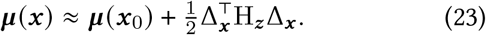

Now, add a first-order approximation 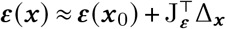 of the noise (posterior uncertainty) to (23), where J_***ε***_ := **∇**_***x***_ ***ε*** (***x***_0_) is the gradient of ***ε*** at ***x***_0_:

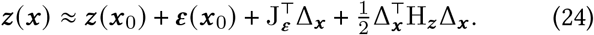

Setting the derivative of (24) in Δ_***x***_ to zero and solving for Δ_***x***_, we find that:

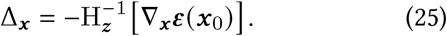

We can construct a covarianc e matrix “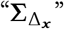” for the location of the peak using (25)

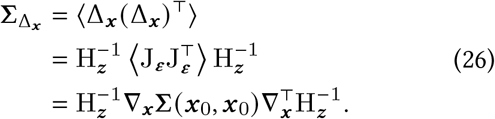

We use the low-rank approximation 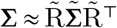 as in (22), where 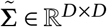 is the low-rank covariance and 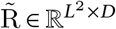 is a semi-orthogonal operator defining our low-rank basis. We can use the Cholesky decomposition to obtain Q ∈ ℝ^*D*×*X*^ such that 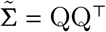 and calculate (26) as

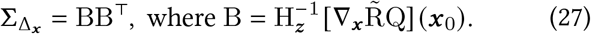

Figure 4d compares field-location confidence intervals obtained either by sampling, or quadratic approximation. These methods agree for well-localized peaks.

### Head-direction dependence

In Figure 5, we show two ways to use LGCP regression to estimate head-direction dependence in grid cells. Firstly, we partitioned the 30-minute recording session into subsets, with sample weights (Fig. 5ab) defined as

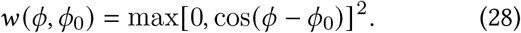

This weighting separates data from opposing head directions (*ϕ*_1_, *ϕ*_2_) = (*ϕ*_0_, *ϕ*_0_+*π*) into non-overlapping subsets.

**Figure 5:**
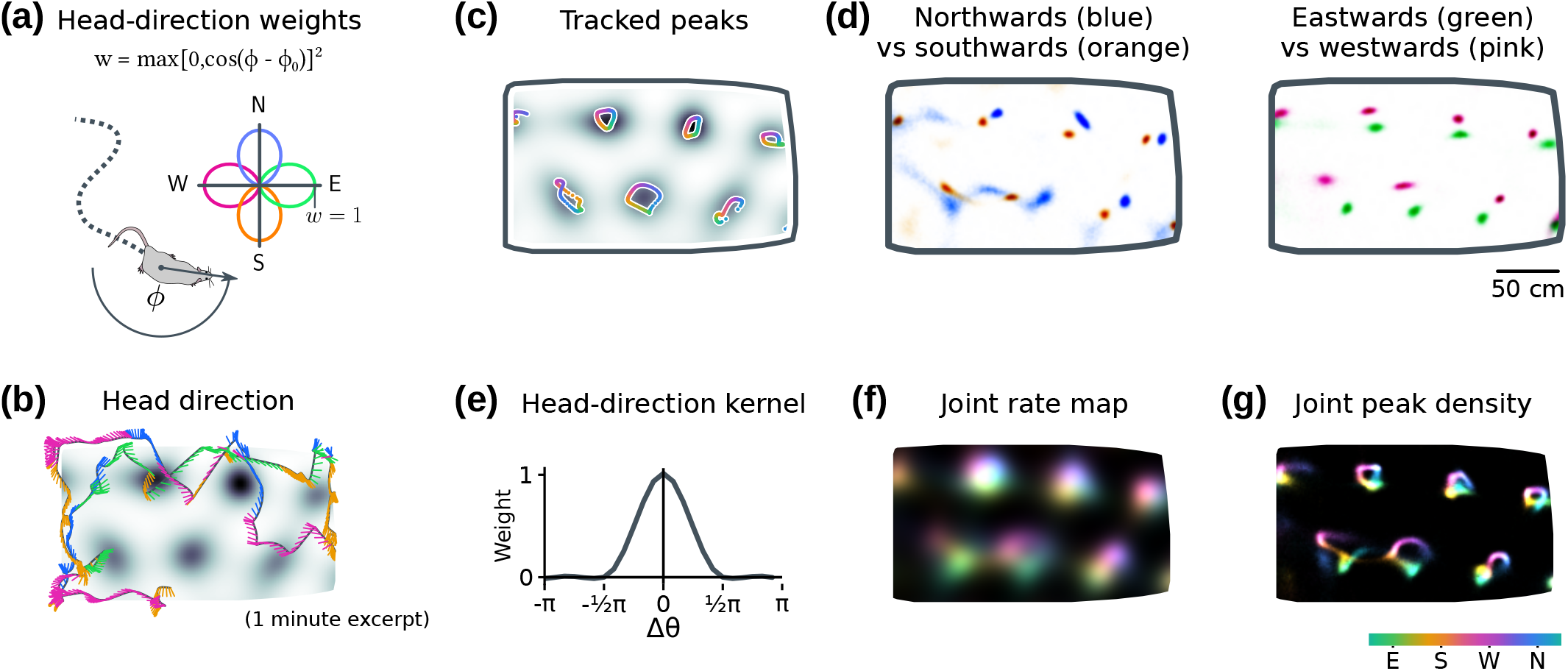
LGCP analysis of joint position–head-direction tuning. We examined head-direction tuning in a cell from Krupic et al. (2018) by conditioning on subsets of the data (a–d), and via estimation of a joint log-rate posterior (e–f) (Methods: *Headdirection analyses*). **(a)** LGCP’s efficiency makes it practical to compare changes in the rate map between subsets of the experimental data. We separated opposing head directions into non-overlapping subsets weighted by cosine similarity between the rat’s head direction and a reference direction. **(b)** The rat’s smoothed head direction, denoted via line segments (colored by nearest cardinal direction) stemming from the smoothed position trajectory (black). **(c)** In this cell, the posterior rate-map peaks depend smoothly on head direction. **(d)** Comparing opposing head directions reveals directionality. **(e)** One can also estimate position and head-direction tuning jointly. Here, we clipped negative Fourier components of the squared-cosine weighting in (a) to form a positive-semidefinite head-direction kernel (shown; 24 direction bins). **(f)** The inferred 2D+heading rate map, depicted in a qualitative color scheme, with preferred head-direction mapped to hue. **(g)** The peaks in the sampled 2D+heading posterior conditioned on each head direction recapitulate the directional shifts seen in Fig. 5c.

Fitting the LGCP estimator over a range of head-direction angles *ϕ*_0_ ∈[0, 2*π*) reveals a continuous and smooth dependence of grid field peaks on head direction (Fig. 5c). Opposing directions (cardinal directions shown in Fig. 5d) show clear differences.

Secondly, we inferred position and head-direction tuning jointly by adding head direction as a third axis to the LGCP regression. To facilitate comparison with Figure 5a–d, we used the same weighting function (28), adjusted to it positive semi-definite “ 𝒦_*ϕ*_ “ (Methods: *Head-direction analyses*; Fig. 5e). The full 2D+direction kernel was a tensor product 𝒦_*ϕ****x***_ = 𝒦_*ϕ*_ ⊗𝒦_***x***_ with a position kernel 𝒦_***x***_ (an optimized version of kernel Fig. 3a-2). The resulting posterior provides a joint distribution of 2D+direction tuning curves, visualized qualitatively in Fig. 5f with head-direction mapped to hue. As in Fig. 4d, one can obtain the distribution of grid-field peaks—in this case conditioned on head direction (Methods: *Head-direction analyses*). This posterior peak-density map (Fig. 5g) recapitulates the head-direction dependence found from applying separate regressions to sub-sampled data (Fig. 5c).

### Estimator performance

We quantify the advantages of LGCP regression over naïve kernel density estimators in Figure 6. We evaluated the estimator performance on a simulated grid map and 30-minute recording session (Fig. 6a) similar to Krupic et al. (2018). On simulated data, the LGCP estimator (optimized grid kernel; Fig. 3a-2) was more accurate than the KDE for a given amount of training data, exhibited less bias than a KDE with bandwidth matching the grid-field scale, and exhibited less variance than a finer-scale KDE (Fig. 6b–d; Methods: *Assessing estimator performance*).

**Figure 6:**
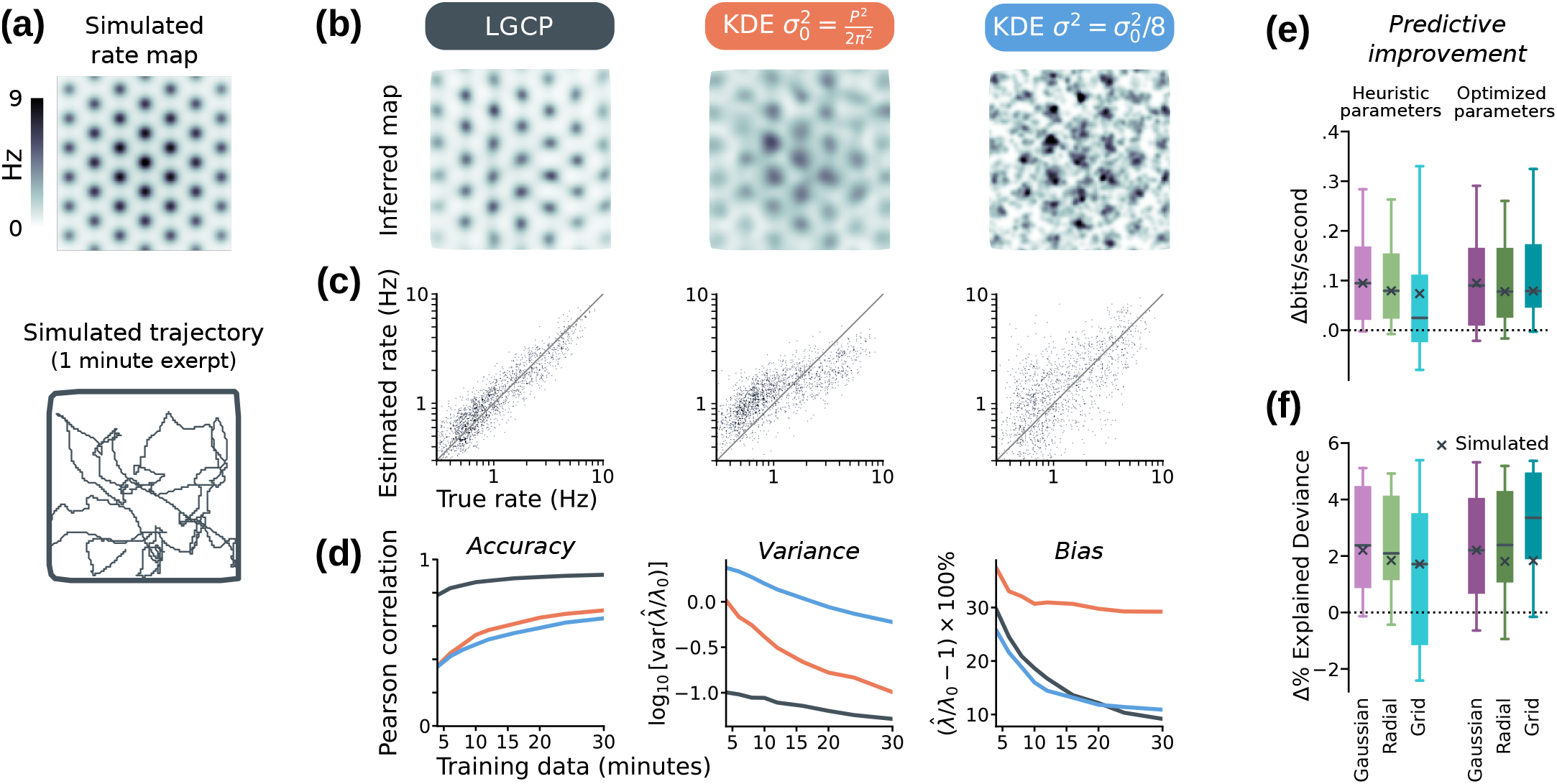
Compared to kernel density estimators, LGCP regression is more efficient and exhibits a superior bias– variance trade-off. **(a)** We sampled Poisson spiking activity from a synthetic grid cell (mean rate 1.2 Hz) throughout 30 minutes of simulated foraging in a square arena. **(b)** Comparison between rate maps recovered via Log-Gaussian Cox Process (LGCP) regression (black, left), a Kernel Density Estimator (KDE) with a kernel width matching the scale of a single grid field (red, middle), and a KDE estimate using a narrower kernel (blue, right). **(c)** The LGCP estimate correlates well with the ground truth, exhibiting less bias than a scale-matched KDE estimator, and less variance than a narrow one. **(d)** Comparison of accuracy (left; Pearson’s correlation between the estimated and recovered rate), variance (middle), and bias (right), between the LGCP and the two KDEs (λ_0_ = true rate, 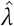 = estimate). **(e**,**f):** We quantified cross-validated (10-fold) estimator performance on 15 randomly selected cells with at least five grid fields from Krupic et al. (2018), and on the simulated data (black ×). We estimated kernel parameters and the posterior rate map, and measured the expected log-likelihood of held-out data. We compared performance to a KDE smoother matched to the grid-field scale (Fig. 6b, middle scenario). **(e)** Cross-validated LGCP expected log-likelihood (adjusted; Methods: *Assessing estimator performance*), relative to KDE log-likelihood baseline. We explored three kernels: A Gaussian kernel (the same one used by the KDE), a radial kernel (kernel Fig. 3a-4), and a grid kernel (kernel Fig. 3a-2). The grid kernel fared worse using heuristic hyperparameters (light bars), but had superior performance once optimized (dark bars). **(f)** The same analysis as (e) reported in terms of % explained deviance (Methods: *Cross-validated performance measures*).

We tested the ability of LGCP regression to predict neuronal activity under cross-validation (Fig. 6e,f). We stress, however, that the application of LGCP regression is not to predict neuronal activity *exactly*, but rather to infer larger-scale features of the grid map by discarding irrelevant fine-scale detail. Nevertheless, the calibrated LGCP estimator consistently matched or exceeded the predictive performance of a kernel-density estimator with bandwidth matched to the grid scale (Methods: *Assessing estimator performance*).

We show two measures of performance in Figure 6e,f: The expected log-likelihood of held-out test data under the regressed LGCP posterior, relative to the log-likelihood of a kernel density estimator (Fig. 6e), and the same results in terms of normalized explained deviance (Methods: *Crossvalidated performance measures*).

## Discussion

We have introduced a variational Bayesian approach to analyzing data from grid cells. We focused on challenges associated with working with grid cells in larger environments, and prioritized computational efficiency to facilitate exploratory analysis of large datasets. Out method incorporates prior assumptions of grid-cell periodicity, is computationally and statistically efficient, yields approximate confidence intervals, and provides way to compare different prior assumptions (i.e. optimize the kernel hyperparameters).

### Caveats

The posterior covariance of a GP regression is only interpretable if the kernel is a true model for the data’s correlation structure. To guard against misspecified kernels, we recommend robust controls for formal hypothesis testing, such as shuffle tests that remove a purported effect form the underlying data. These are computationally intensive but feasible, requiring resources similar to shuffle controls for a large GLM regressions.

We have assumed that a single kernel captures the correlation structure at all locations, whereas grid cells are known to display subtle changes e.g. near boundaries (Hägglund et al., 2019). While it is possible to relax the assumptions encoded in the kernel (e.g. by summing multiple angular/radial kernels to create orientation/period flexibility), a spatially varying solution may be preferable. Research into non-stationary (or spatially inhomogeneous) kernels is ongoing (e.g. Paun et al. 2023), but we are aware of no suitable advances for the specific problem. In principle, one could merge maps from different priors in different regions of the arena *post hoc*. We leave such explorations for the future.

### Generality and applicability

The GP estimators described here combine aspects of traditional kernel-density smothers and Poisson generalized linear models with a principled (periodic) prior. The advantages of GP regression over the KDE are that GP regression (1) adapts to smooth more where data are limited (2) can be used to infer parameters of the spatial correlation structure (e.g. identify grid period), and (3) can employ priors with nearest-neighbor interactions, allowing the smoothing radius to exceed the size of a single grid field.

Our implementation is broadly applicable to problems that are large enough for the cubic complexity of naïve GP estimators to become burdensome, but which are unsuitable for existing sparse or low-rank algorithms. It can be applied to intermediate-size spatiotemporal inference problems with kernels that are sparse in the frequency domain.

Although we focused on the Poisson observation model, our approach generalizes to other observation models in the natural exponential family. The main caveat is other observation models may lack a convenient closed-form expression for the expected firing rate, and these terms may need to be approximated via sampling.

### Future work

The methods we have described are suitable as a drop-in replacement for the smoothing and kernel density estimators currently used to analyze grid cell activity. In addition to providing a principled smoother that is aware of a grid cell’s spatial scale, our approach provides approximate confidence intervals. These algorithms have considerable potential, and could be extended e.g. to incorporate tuning to additional behavioral covariates, or additional latent rate fluctuations that are correlated in time.

We plan to apply these methods in our own work. We hope that others will as well, in addition to building upon these approaches to design new algorithms for spatiotemporal statistics in the study of spatial navigation.

## Code and data availability

We have provided a reference implementation in Python online at Github (http://github.com/michaelerule/lgcpspatial). We have included the fifteen test cells from Krupic et al. (2018) required to reproduce the figures and demonstrations in this manuscript. Any use of these data should cite Krupic et al. (2018).

## Acknowledgments

M.E.R. is supported by a Leverhulme and Isaac Newton Trust fellowship ECF-2020-352. P.C.V is supported by MRC, Frank Elmore Fund, and University of Cambridge School of Clinical Medicine. J.K. is a Wellcome Trust/Royal Society Sir Henry Dale Fellow (206682/Z/17/Z) and is supported by Dementia Research Institute (DRICAMKRUPIC18/19), Isaac Newton Trust/Wellcome Trust ISSF/University of Cambridge Joint Research Grant, Kavli Foundation Dream Team project (RG93383), Isaac Newton Trust [17.37(t)], and NVIDIA Corporation. T.O’L. is supported by ERC grant 716643 FLEXNEURO and HFSP grant RGY0069/2017.

## Methods

### Finding the posterior mode

One approach to estimating the firing rate map is to find the ***z*** that maximizes the log-posterior (7). This is the *Maximum a Posteriori* (MAP) estimator, and can be solved via gradient ascent. The gradient of (7) in ***z*** is:

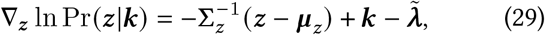

where we have defined 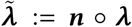 as the vector of estimated firing-rates weighted by the number of visits to each location ***n*** (° denotes element-wise multiplication).

The (negative) log-posterior (7) is convex, and well approximated as locally-quadratic. The Newton-Raphson method is applicable, and much faster than gradient descent. This uses the curvature (Hessian, “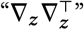”) of (7) in ***z***:

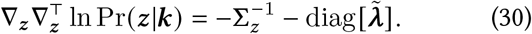

On each iteration, Newton-Raphson must solve the update

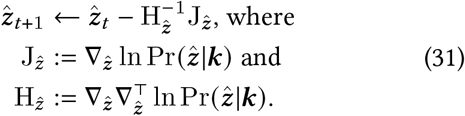

The Hessian and Jacobian for the MAP estimator are the same as those for the variational posterior mean (17), but with the expected rate 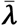 replaced by the point estimate 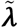. To compute 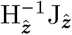 quickly in high dimensions, we used an inexact Newton-Raphson method (Dembo et al., 1982) that approximates 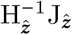 each iteration of (31) via a preconditioned Krylov method (Methods: *Newton-Krylov methods*)

### Newton-Krylov methods

Naïve algorithms for multiplying or inverting dense matrices have cubic complexity, making expressions such as (31) computationally prohibitive in higher dimensions (c.f. Liu et al., 2020). Thankfully, modern Krylov-subspace algorithms for solving large linear systems ***A***^−1^***v*** only require a function that can compute the matrix-vector-product ***u*** ↦***Au***. This can be computed quickly if our matrix ***A*** has special structure (Chan and Jackson, 1984; Brown and Saad, 1990; see Knoll and Keyes, 2004 for review). Fast solutions exist if the covariance kernel (or its inverse) is sparse (e.g. Luttinen and Ilin, 2009; Kiiveri and De Hoog, 2012; Gal et al., 2014; Cseke et al., 2016) or Toeplitz/circulant (eg. Jensen et al., 2021).

Algorithms combining Newton-Raphson iteration with Krylov methods were first developed for very large, sparse, Gaussian process models (Kiiveri and De Hoog, 2012; Cseke et al., 2016), but apply to any problem where ***Au*** can be computed quickly but ***A***^−1^***u*** is impractical. In our case, our prior covariance **Σ**_*z*_ is a convolution, and we calculate 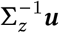 quickly as point-wise multiplication in in the spatial-frequency domain (Methods: *Working in a lowrank subspace*). We can therefore calculate the Hessianvector product 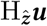 quickly, which allows us to expediently calculate 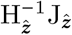 using a Krylov-subspace solver.

In our tests, we found that Scipy’s (Virtanen et al., 2020) implementation of the minimum residual Krylov-subspace algorithm (MINRES; Paige and Saunders, 1975) provided the best balance of speed and stability. To make complex-valued spatial-frequency components compatible with Krylov solvers designed for real-valued matrices, we used the Hartley transform rather than the Fourier transform (Methods: *Working in a low-rank subspace*).

Krylov-subspace algorithms benefit from a preconditioner “*M*” that approximates the inverse (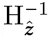 in our case). We used the prior covariance kernel for this, *M* = **Σ**_*z*_, also computed as a convolution via pointwise multiplication in a low-rank spatial-frequency domain (see Algorithm 1).

### Connection to the Laplace Approximation

The Laplace approximation (Fig. 2b) models the posterior uncertainty in the MAP-estimated log-rate 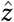 as a Gaussian centered at 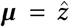, and with the covariance equal to the negative-inverse of the Hessian (30) evaluated at 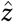:

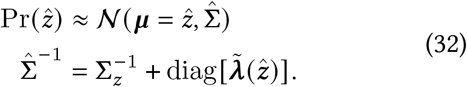

Intuitively, (32) says that the Laplace approximation models the posterior precision as a sum of the prior precision and a diagonal matrix representing information from spiking observations.

Note the similarity between the derivatives of the variational mean (17) and those of the MAP estimator (29) and (30), and the similarity between the Laplace-approximated posterior variance (32) and the covariance update for variational Bayes (19) and (45). Optimizing the variational mean is tantamount to calculating the MAP estimator using the expected rate ⟨***λ***⟩_**z**_ rather than a point estimate ***λ***.

Likewise, updating the posterior covariance is tantamount to applying the Laplace approximation that the variational mean, again using the expected rate rather than a point estimate.

### Derivatives

In this section, we derive the gradients of the evidence lower bound (15) with respect to **Σ** and ***q*** (Equations (18) and (20) in the main text, respectively).

First, we obtain the derivative of the evidence lower bound (15) with respect to the posterior covariance matrix **Σ**. Consider the derivative of the term term ***n***^⊤^ ⟨***λ***⟩ with respect to individual elements **Σ**_*i j*_. We use Einstein summation notation, wherein sums over repeated indecies is implied:

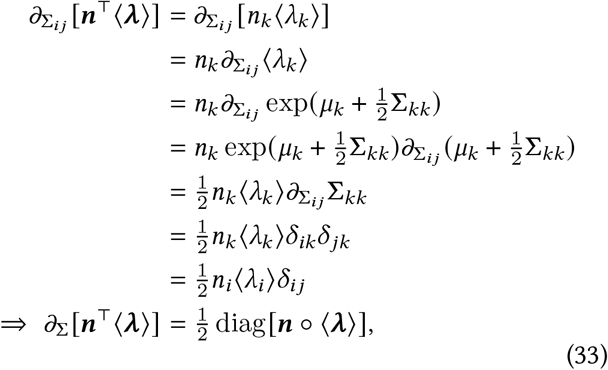

where *δ*_ab_ is the Kronecker delta (1 if a = b and 0 otherwise).

The derivative of the term 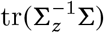 in (15) is given by identity (100) in The Matrix Cookbook (Petersen et al., 2008), and is 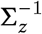. The derivative of the term ln |**Σ**| is given by identity (57) in The Matrix Cookbook (Petersen et al., 2008), (assuming |**Σ**| ≠ 0), and is **Σ**^−1^. Overall, then, the derivative of (15) in **Σ** is:

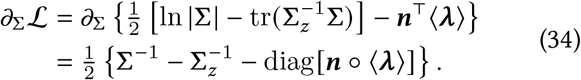

Let 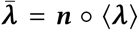. For the parameterization 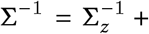+ diag[***q***], (34) simplifies to:

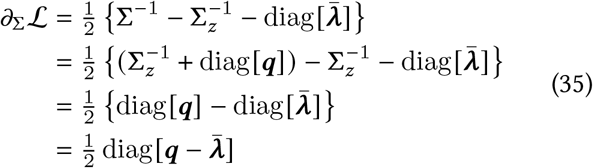

We can now obtain the derivative of the evidence lower bound (15) in ***q*** via the chain rule, 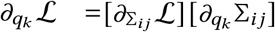. To obtain *∂*_*q*k_ **Σ**_*i j*_, we will need identity (59) in The Matrix Cookbook (Petersen et al., 2008), which provides the chain rule for the derivative of the inverse of a matrix, *∂*_*x*_ [***A***(*x*)^−1^] = −***A***^−1^ [*∂*_*x*_ ***A***]***A***^−1^. We will let A=Σ^-1^:

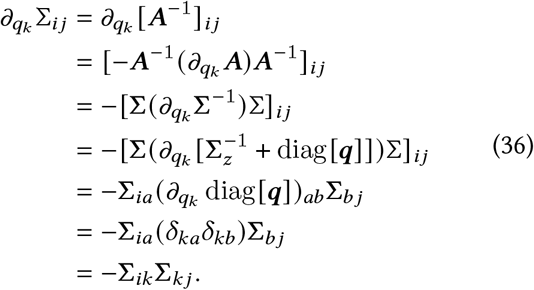

Combining (35) and (36) gives:

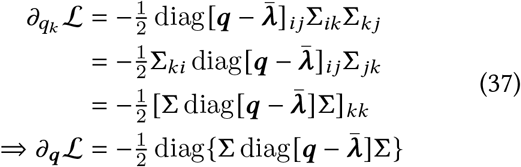

### Working in a low-rank subspace

Working in a lowrank subspace can make large problems tractable. We first find a low-rank approximation of the prior covariance **Σ**_*z*_, and then perform inference within this subspace.

The prior covariance **Σ**_*z*_ is defined by a convolution kernel. The components of this kernel “***ξ*** “ in spatial-frequency (Fourier) space are the eigenvalues of **Σ**_*z*_:

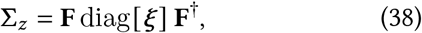

where **F** is the unitary Fourier transform and † denotes the conjugate (Hermitian) transpose.

In practice, many spatial frequency components will be close to zero. These are frequencies where the prior assigns very little probability. We work in a low-rank space consisting only of those directions in **Σ**_*z*_ where the prior has assigned non-negligible variance. We retain the *D*≤ *L*^2^ components “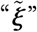” whose magnitude in the prior covariance kernel is at least 10% of the eigenvalue of the prior covariance with the largest magnitude “*ξ*_max_”:

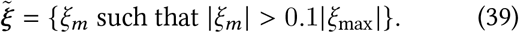

The low-rank approximation to the posterior covariance can then be calculated as

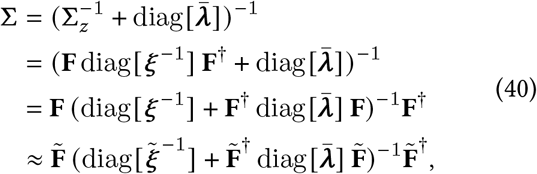

where 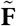 is the (unitary) Fourier transform retaining only the non-negligible components 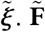. is not invertable, but since it is semi-orthogonal, 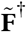 is its pseudoinverse.

Note that the Kullback-Leibler divergence contribution to the evidence lower bound (13) contains a constant factor 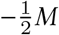 that depends on the number of dimensions *M* = *L*^2^ in the our multivariate-Gaussian prior. When working in a low-rank *D* < *M* subspace, this term should be replaced by 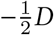 to ensure that the evidence lower-bound can be compared between models with low-rank subspaces of different ranks.

Fourier coefficients can take on complex values. This creates compatibility and performance issues with standard numerical linear algebra software. To address this, we use a real-valued relative of the Fourier transform called the Hartley transform (Hartley, 1942).

We denote the Hartley transform as **R**, and the transform with negligible frequencies discarded as 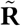. The Hartley transform is calculated by summing the real and imaginary components of the Fourier transform

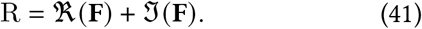

If **F** is the unitary Fourier transform, then R is also unitary.

Equations in (38)-(43) work similarly with the Hartley transform, replacing **F** with **R**, and replacing Hermitian transposes with ordinary transposes. Since the (circulant) prior **Σ**_*z*_ is symmetric, its Fourier coefficients ξ are real-valued, and the Hartley-transform coefficients for the prior are identical to the Fourier coefficients. We can write (40) using the Hartley transform as

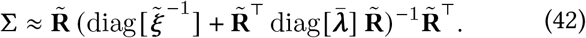

We denote this approximation as 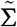. During inference, only *N < M* bins with nonzero observations (*n*_*m*_ *>* 0) contribute to the expected log-likelihood, and we can further truncate 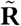 to an *N* × *D* matrix for efficiency.

The frequency-subspace representation simplifies some of the matrix calculations. Let *L* be the size of the environment, and *L*^2^ be the total number of spatial bins. A *L* ×*L* array can be converted into frequency space using the 2D fast Fourier transform, which costs **𝒪** (*L*^2^ log (*L*)). If we retain only *D* components, the relevant transform has dimensions *L*^2^*D* and the cost is **𝒪** (*L*^2^*D*). Ordinary matrix multiplication can outperform the FFT when *D*∼ **𝒪** (log (*L*)). Since grid cells display only a narrow range of spatial scales, *D* can be small, and the complexity of each optimization iteration is competitive with simpler estimators.

We perform most calculations in this low-rank space, and never explicitly construct the posterior covariance. The only calculation which cannot be performed in the low-rank space is the calculation of the expected firing-rates at each location, 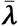, which we address in *Methods: Calculating the expected firing rate*. This has complexity **𝒪** (*D*^2^*N* +*D*^3^), where *N* is the number of spatial bins containing observations.

When operating in a low-rank subspace, it is important that the mean of the variational posterior also be expressed in this subspace. Leaving ***μ*** in the full-rank space creates a poorly-conditioned problem, since several directions will be ignored when calculating gradients using low-rank approximations.

### Calculating the expected firing rate

For Gaussian ***z*** ∼ **𝒪** (***μ*, Σ**) and exponential firing-rate nonlinearity, the firing rate ***λ*** = exp (***z***) is log-normally distributed. The mean of this distribution, ⟨*λ*⟩, has the closed-form expression 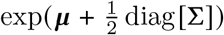(c.f. Rule and Sanguinetti, 2018). Evaluating this expression requires the diagonal of the posterior covariance matrix.

In the low-rank subspace (42), these diagonal elements **Σ**_*ii*_ can be calculated with the following procedure:

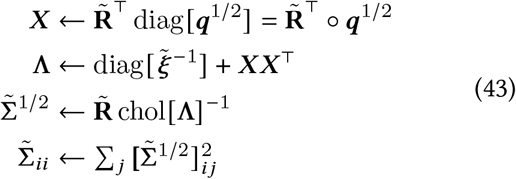

In (43), we first project the (square root of the) precision update diag [***q***^1/2^] into the low-rank subspace. We then obtain the inverse posterior covariance in the lowrank space, “Λ”. Rather than invert this directly, we compute its Cholesky factorization and use a triangular inverse solver. We expand this from the low-rank subspace using the inverse FFT. This provides a factor 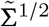 of the low-rank approximation to the posterior covariance such that 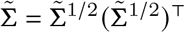, from which we extract the diagonal variances. This factor is also useful for sampling from the variational posterior (22).

### Iteratively estimating *q*

From (20), we see that ***q*** must equal 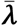 to maximize the evidence lower bound (15) (for fixed ***μ***). However, since 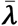 depends on ***q***, this must be solved self consistently. This can be solved by ascending the simpler gradient 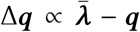. Taking discrete steps yields the following fixed-point iteration:

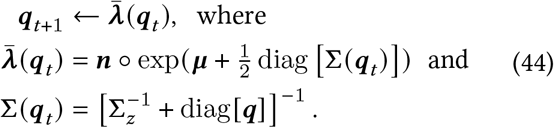

In practice, we implement this by iterating the marginal posterior variances ***v*** = diag [**Σ**]. This is amounts to a different parameterization of (44):

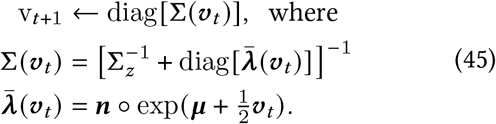

Challis and Barber (2013) note that the iteration in (44) may diverge. In practice, we have found that the reparameterized iteration in (45) always converges when starting from ***v*** = 0, provided one re-optimizes the posterior mean before each step according to (17), and provided the prior is sufficiently well-conditioned. Note that **Σ** (***v***_*t*_) remains bounded in the parameterization in (45) as 0 ≼ **Σ** (***v***_*t*_) ≼ Σ_z_. This implies that the iteration cannot diverge to infinity if the prior precision 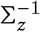 is full rank (≼ is the Loewner order of positive semidefinite matrices). Since these iterations are simply gradient descent with a step size of 1, instability can be remedied by choosing a smaller step-size, if encountered.

### Binning data

For clarity, we presented the derivations in this manuscript in terms of piecewise-constant spatial basis functions. In practice, linearly interpolated binning provides better resolution for a given grid size, and this is what we used in the provided reference implementation.

For each visit and/or spike at location location ***x***, we distributed the point mass at ***x*** over a 2 ×2 neighborhood of adjacent bins via linear interpolation. This amounts to using square-pyramidal basis functions to provide a piecewise-linear model the inferred firing-rate map (compare to Figure 1 in Cseke et al., 2016).

Since most calculations are performed directly on the spatial-frequency components of the grid map, choices for spatial binning only affect the numeric integration of the data likelihood (5) over the spatial domain. Locally-constant binning amounts to using the Riemann sum to compute this integral, and linearly interpolated binning amounts to using the trapezoid rule. Linear interpolation improves resolution compared to a piecewise-constant model, but conceptually there are no substantive differences.

### Initializing parameters

#### Grid period P and orientation

We estimated the grid period *P* using the radial autocorrelogram of the firingrate histogram ***y*** = ***k*** /***n***, calculated by averaging the 2D spatial autocorrelogram over all angles. The radial autocorrelogram “***R***_*ρ*_ “ for a period-*P* periodic spatial signal is given by Equation (9):

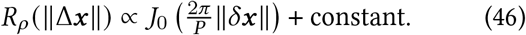

The location “Δ_p_ “ of the first nonzero peak of ***R***_*ρ*_ (∥Δ***x***∥) depends on *P*, and we can solve for *P* given Δ_p_ as

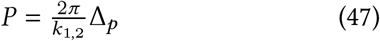

where *k*_1,2_ is the second zero of the first-order Bessel function of the first kind.

When using the oriented kernel (Fig. 3a-2), we estimated the grid orientation based on the phase of a 6-fold periodic sinusoid fit to the spatial autocorrelogram at distance *r* = *P*/(2*πk*_1,2_).

#### *Heuristic μ and prior mean μ*_z_

We used a Gaussian kernel density smoother to estimate foreground 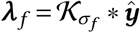 and background 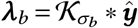 rate maps (*σ*_b_ = *P π* ; *σ*_b_ = 5*σ*_b_). We use this background log-rate map for the prior mean ***μ***_**z**_ in variational inference. We used the foreground as an initial guess when optimizing the posterior mean.

#### *Kernel height* 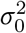 ***and constant offset*** *c*

We calculated an initial estimate of the log-rate as the difference between the log-foreground and log-background maps. The variance of this map was then used to initialize the kernel height, 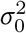. The kernel’s constant offset *c* controls how confident we are in our prior assumptions about the average log-firing-rate across the environment. The average log-rate is the average (“DC”) component of the prior mean ***μ***_**z**_. We set *c* = 10^3^ to leave the inference procedure free to adjust the mean log-rate.

***Grid search:*** For the analyses shown in this paper, we refined kernel hyperparameters in a two-step process. Starting from heuristically initialized parameters, we estimated, 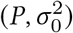)via grid-search with an orientation-agnostic kernel (Fig. 3a-4). We recursively searched nearby values of **Θ** until we found a local maximum. We re-used solutions for the parameters of the variational posterior from previous choices of **Θ** as initial guesses for optimizing new **Θ** to reduce computational cost. Then, we identified orientation *θ*_0_ by leaving 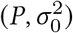) fixed an sweeping a range of angles in [0, *π*/3), Finally, we re-optimized 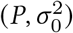) for a grid kernel with orientation *θ*_0_ (Fig. 3a-2).

### Head-direction analyses

For Figure 5, head direction was tracked via a head-mounted infrared LED (see Krupic et al., 2018 for details). We converted the recorded head direction *ϕ*_raw_ (*t*) into cosine and sine components, {*ϕ*_*x*_, *ϕ*_*y*_ }= {cos *ϕ*_raw_ (*t*), sin *ϕ*_raw_ (*t*) }. We then imputed missing data via linear interpolation and smoothed {*ϕ*_*x*_, *ϕ*_*y*_ } with a 2 Hz low-pass Savitsky-Golay filter, yielding smoothed estimates 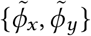 and head direction 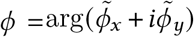. We used data from the entire experimental session to optimize the period, variance, and direction of a local-neighborhood grid kernel (kernel Fig. 3a-2) via grid search. We used this spatial kernel “ **𝒦**_***x***_ “ for all subsequent regressions.

Analyzing head-direction via weighted subsets of the data reduces to the 2D inference problem. For each reference direction *ϕ*_0_, we defined a weighting function *w* (*t*) ∈[0, 1] as *wt* = max[0, cos(*ϕt* − *ϕ*0)]2 L(Eq. (28)). We calculated weighted visit counts 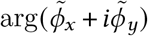 and spike counts 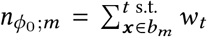 (compared to Equation (4)). Inference of a heading-conditioned rate map amounts to inferring a 2D position rate map using these weighted counts.

To construct joint 2D+direction LGCP regression (Fig. 5e– g), one treats the time-varying head direction as a third spatial dimension. The only difference from the spatial case is that the head-direction axis does not require padding to avoid circular wrap-around. We defined a grid of *D* = 24 head directions uniformly spaced around the circle, and binned the smoothed head direction using linear interpolation (Methods: *Binning data*).

To facilitate comparison between approaches, we modified the weighting function (28) into a positive semidefinite kernel by clipping its negative eigenvalues to zero, i.e. ***𝒦***_*ϕ*_ (*ϕ, ϕ*^′^) = F^−1^ max 0, [F {*w* (*ϕ, ϕ*^′^)}]. We constructed the joint kernel ***𝒦***_*ϕ****x***_ = ***𝒦***_*ϕ*_ ⊗***𝒦***_***x***_ as a Kronecker product in the spatial domain, then discarded all but the *D* = 1000 largest components Fourier domain to generate a lowrank subspace. Inference of the posterior log-rate density is then identical to the 2D case. Unlike Savin and Tkacik (2016), we do not use the Kronecker structure of the (direction ⊗ position) prior in the inference, but rather infer the joint posterior in a low-rank subspace.

We calculated the head-direction-dependent peak-density map in Fig. 5g by drawing 2D+direction samples from the inferred posterior distribution, conditioning on each head-direction separately, and identifying local maxima with a radius of *P* /2.5 of the grid period *P* (with peak locations up-sampled via quadratic interpolation).

### Assessing estimator performance

In Figure 6, we assess the LGCP estimator performance on both simulated and experimental data. We simulated an ideal grid cell on a 90 ×90 grid with log-rate as in (8), scaled to a mean rate of 1.2 Hz, and with a period of 13 bins. For comparison to the experimental results throughout the text, if each bin were 2 ×2 cm^2^, this would correspond to a 1.8 ×1.8 meter enclosure and a cell with a period of 26 cm. We simulated 30 minutes of random exploration at 50 samples per second as Brownian motion (*σ*^2^ = 0.02 bin^2^/second) clipped to the arena boundaries, filtered twice with a first-order exponential smoother (**τ** = 190 ms).

We compared the accuracy, bias, and variance of the LGCP and Kernel Density Estimator (KDE) in Fig. 6b–d. We defined a “scale-matched” Gaussian KDE with variance 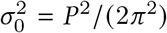. This matches the curvature of the Gaussian kernel at Δ***x*** = 0 with that of the radial autocorrelation (9), yielding a Gaussian kernel that approximates the size and shape of a single grid field. We also defined a “finer-scale” KDE kernel, with variance 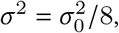, which was more noisy, but provided a less biased estimate in expectation. To assess accuracy, bias, and variance as a function of data size (i.e recording length), we partitioned the synthetic data into 15 blocks and sampled bootstrapped training datasets of varying duration, with replacement (200 samples). We kept the kernel parameters fixed (for all estimators), rather than re-estimating them on each sample (see Methods: *Cross-validated performance measures* for an assessment that incorporates hyperparameter uncertainty).

### Cross-validated performance measures

We compared the ability of the LGCP and scale-matched 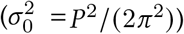 KDE to predict spiking activity on held-out test data in Figure 6e,f. We used a simulated dataset and fifteen randomly chosen cells from Krupic et al. (2018) (of those with at least five grid fields).

We compared three different kernels for the LGCP estimator: (i) A Gaussian Radial Basis Function (RBF), with variance 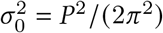 identical to that of the KDE; (ii) A radial kernel (Figure 3a-4), which included no assumptions about grid orientation, and (iii) A local-neighborhood grid kernel (Figure 3a-2). The Gaussian kernel provides a fair comparison with the KDE, and the radial vs. grid kernel performance emphasizes the importance of hyperparameter optimization.

We assessed performance under 10-fold cross-validation. We tested both heuristic (Methods: *Initializing parameters*) and grid-search-optimized kernel hyperparameters. Hyperparameter estimates were repeated for each block with held-out data excluded. The reference KDE bandwidth was fixed at the cells “true” period as identified by the optimal kernel parameters on the whole dataset.

We assessed LGCP performance using the expected loglikelihood (14) of the held-out test data under the inferred posterior distribution (or, for the KDE: the point estimate (5)). Since changes in mean-rate between train/test data are uninteresting for inferring spatial variations in tuning, we adjusted the predicted mean-rate to match the test data before evaluating the (expected) log-likelihood (“adjusted log-likelihood”). We also report performance in terms of change in the % explained deviance, in analogy to a normalized ***R***^2^ statistic from linear regression. We defined the “null” model as one that simply guesses the mean-rate on the test data 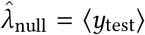 (worst-case performance), and the “saturated” model as 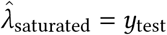 (theoretical maximum of the Poisson likelihood). Normalized explained deviance is given as

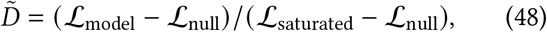

where ℒare the (expected) log-likelihoods of the respective models. We report the improvement in (48) relative to the KDE baseline (×100%) in Figure 6f.

#### Algorithm 1

*Iterative procedure for variational-Bayesian log-Gaussian Cox process regression;* ° denotes element-wise vector and matrix products, with ***u*** ° A:= diag[***u***]A and A ° ***u***:=A diag[***u***], and (·)^°·^ denotes element-wise power.

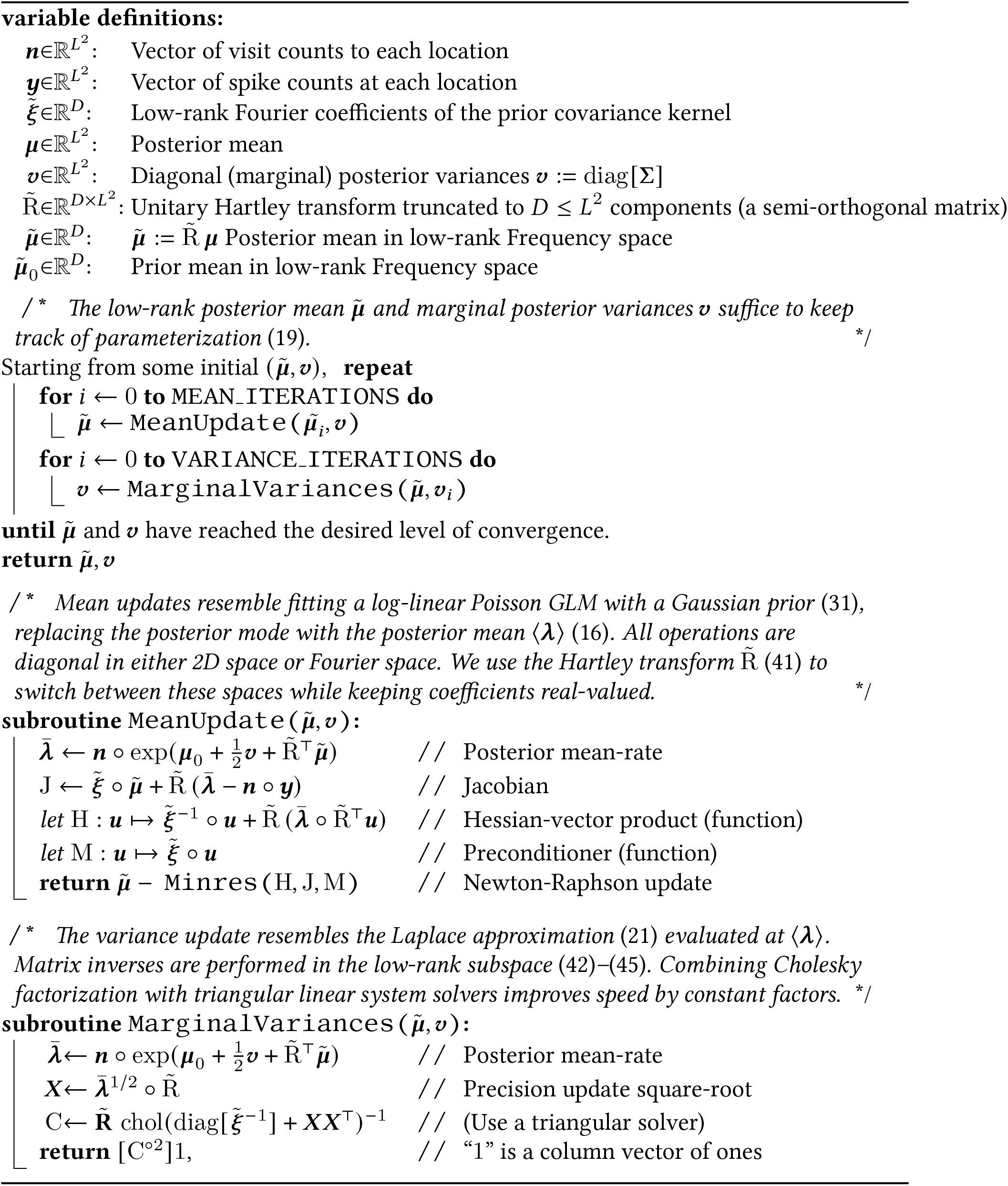

## References

Bradbury, J., Frostig, R., Hawkins, P., Johnson, M. J., Leary, C., Maclaurin, D., & Wanderman-Milne, S. (2018). JAX: composable transformations of Python+NumPy programs http://github.com/google/jax.

Brandman, D. M., Burkhart, M. C., Kelemen, J., Franco, B., Harrison, M. T., & Hochberg, L. R. (2018). Robust closed-loop control of a cursor in a person with tetraplegia using Gaussian process regression. Neural computation, 30(11), 2986–3008, https://doi.org/10.1162/neco_a_01129.

Brandon, M. P., Bogaard, A. R., Libby, C. P., Connerney, M. A., Gupta, K., & Hasselmo, M. E. (2011). Reduction of theta rhythm dissociates grid cell spatial periodicity from directional tuning. Science, 332(6029), 595–599, https://doi.org/10.1126/science.1201652.

Brown, P. N. & Saad, Y. (1990). Hybrid Krylov methods for nonlinear systems of equations. SIAM Journal on Scientific and Statistical Computing, 11(3), 450–481, https://doi.org/10.1137/0911026.

Challis, E. & Barber, D. (2013). Gaussian Kullback-Leibler approximate inference. Journal of Machine Learning Research, 14(8) http://www.jmlr.org/papers/v14/challis13a.html.

Chan, T. F. & Jackson, K. R. (1984). Nonlinearly preconditioned Krylov subspace methods for discrete Newton algorithms. SIAM Journal on scientific and statistical computing, 5(3), 533–542, https://doi.org/10.1137/0905039.

Chaudhuri-Vayalambrone, P., et al. (2023). Simultaneous representation of multiple time horizons by entorhinal grid cells and ca1 place cells. Cell Reports, 42(7), https://doi.org/10.1016/j.celrep.2023.112716.

Cox, D. R. (1955). Some statistical methods connected with series of events. Journal of the Royal Statistical Society: Series B (Methodological), 17(2), 129–157, https://doi.org/10.1111/j.2517-6161.1955.tb00188.x.

Cseke, B., Zammit-Mangion, A., Heskes, T., & Sanguinetti, G. (2016). Sparse approximate inference for spatiotemporal point process models. Journal of the American Statistical Association, 111(516), 1746–1763, https://doi.org/10.1080/01621459.2015.1115357.

Dembo, R. S., Eisenstat, S. C., & Steihaug, T. (1982). Inexact Newton methods. SIAM Journal on Numerical analysis, 19(2), 400–408, https://doi.org/10.1137/0719025.

Duncker, L. & Sahani, M. (2018). Temporal alignment and latent Gaussian process factor inference in population spike trains. bioRxiv, (pp. 331751)., https://doi.org/http://proceedings.neurips.cc/paper/2018/hash/d1ff1ec86b62cd5f3903ff19c3a326b2-Abstract.html.

Frigola, R., Chen, Y., & Rasmussen, C. E. (2014). Variational Gaussian process state-space models. In Advances in neural information processing systems (pp. 3680–3688). http://proceedings.neurips.cc/paper/2014/hash/139f0874f2ded2e41b0393c4ac5644f7-Abstract.html.

Gal, Y., van der Wilk, M., & Rasmussen, C. E. (2014). Distributed variational inference in sparse gaussian process regression and latent variable models. https://doi.org/10.48550/arXiv.1402.1389.

Gerlei, K., Passlack, J., Hawes, I., Vandrey, B., Stevens, H., Papastathopoulos, I., & Nolan, M. F. (2020). Grid cells are modulated by local head direction. Nature communications, 11(1), 1–14, https://doi.org/10.1038/s41467-020-17500-1.

Ginosar, G., Aljadeff, J., Burak, Y., Sompolinsky, H., Las, L., & Ulanovsky, N. (2021). Locally ordered representation of 3d space in the entorhinal cortex. Nature, 596(7872), 404–409, https://doi.org/10.1038/s41586-021-03783-x.

Hafting, T., Fyhn, M., Molden, S., Moser, M.-B., & Moser, E. I. (2005). Microstructure of a spatial map in the entorhinal cortex. Nature, 436(7052), 801–806, https://doi.org/10.1038/nature0372.

Hagglund, M., Mørreaunet, M., Moser, M.-B., & Moser, E. I. (2019). Grid-cell distortion along geometric borders. Current Biology, 29(6), 1047–1054, https://doi.org/10.1016/j.cub.2019.01.074.

Hartley, R. V. (1942). A more symmetrical fourier analysis applied to transmission problems. Proceedings of the IRE, 30(3), 144–150, https://doi.org/10.1109/JRPROC.1942.234333.

Jensen, K., Kao, T.-C., Tripodi, M., & Hennequin, G. (2020). Manifold GPLVMs for discovering non-euclidean latent structure in neural data. Advances in Neural Information Processing Systems, 33, 22580–22592 http://proceedings.neurips.cc/paper/2020/hash/fedc604da8b0f9af74b6cfc0fab2163c-Abstract.html.

Jensen, K. T., Kao, T.-C., Stone, J. T., & Hennequin, G. (2021). Scalable Bayesian GPFA with automatic relevance determination and discrete noise models. bioRxiv http://proceedings.neurips.cc/paper/2021/hash/58238e9ae2dd305d79c2ebc8c1883422-Abstract.html.

Keeley, S. & Pillow, J. (2018). Introduction to Gaussian processes http://pillowlab.princeton.edu/teaching/statneuro2018/slides/notes12_GPs.pdf.

Keeley, S., Zoltowski, D., Yu, Y., Smith, S., & Pillow, J. (2020). Efficient non-conjugate Gaussian process factor models for spike count data using polynomial approximations. In International Conference on Machine Learning (pp. 5177–5186).: PMLR http://proceedings.mlr.press/v119/keeley20a.html.

Keinath, A. T., Epstein, R. A., & Balasubramanian, V. (2018). Environmental deformations dynamically shift the grid cell spatial metric. Elife, 7, e38169, https://doi.org/10.7554/eLife.38169.

Kiiveri, H. & De Hoog, F. (2012). Fitting very large sparse Gaussian graphical models. Computational Statistics & Data Analysis, 56(9), 2626–2636, https://doi.org/10.1016/j.csda.2012.02.007.

Killian, N. J., Jutras, M. J., & Buffalo, E. A. (2012). A map of visual space in the primate entorhinal cortex. Nature, 491(7426), 761–764, https://doi.org/10.1038/nature11587.

Knoll, D. A. & Keyes, D. E. (2004). Jacobian-free Newton– Krylov methods: a survey of approaches and applications. Journal of Computational Physics, 193(2), 357–397, https://doi.org/10.1016/j.jcp.2003.08.010.

Krupic, J., Bauza, M., Burton, S., & O’Keefe, J. (2018). Local transformations of the hippocampal cognitive map. Science, 359(6380), 1143–1146, https://doi.org/10.1126/science.aao496.

Langston, R. F., et al. (2010). Development of the spatial representation system in the rat. Science, 328(5985), 1576–1580, https://doi.org/10.1126/science.1188210.

Liu, H., Ong, Y.-S., Shen, X., & Cai, J. (2020). When Gaussian process meets big data: A review of scalable GPs. IEEE transactions on neural networks and learning systems, 31(11), 4405–4423, https://doi.org/10.1109/TNNLS.2019.2957109.

Luttinen, J. & Ilin, A. (2009). Variational Gaussianprocess factor analysis for modeling spatiotemporal data. Advances in neural information processing systems, 22, 1177–1185 http://proceedings.neurips.cc/paper/2009/hash/4a47d2983c8bd392b120b627e0e1cab4-Abstract.html.

MacKay, D. J. (1998). Introduction to Gaussian processes. NATO ASI series F computer and systems sciences, 168, 133–166.

Paige, C. C. & Saunders, M. A. (1975). Solution of sparse indefinite systems of linear equations. SIAM journal on numerical analysis, 12(4), 617–629, https://doi.org/10.1137/0712047.

Paninski, L. (2004). Maximum likelihood estimation of cascade point-process neural encoding models. Network: Computation in Neural Systems, 15(4), 243, https://doi.org/10.1088/0954-898X 15 4 002.

Park, M., Weller, J. P., Horwitz, G. D., & Pillow, J. W. (2014). Bayesian active learning of neural firing rate maps with transformed Gaussian process priors. Neural computation, 26(8), 1519–1541, https://doi.org/10.1162/NECO_a_00615.

Paun, I., Husmeier, D., & Torney, C. J. (2023). Stochastic variational inference for scalable non-stationary gaussian process regression. Statistics and Computing, https://doi.org/10.1007/s11222-023-10210-w.

Petersen, K. B., Pedersen, M. S., et al. (2008). The matrix cookbook. Technical University of Denmark, 7(15), 510.

Quinonero-Candela, J. & Rasmussen, C. E. (2005). A unifying view of sparse approximate Gaussian process regression. The Journal of Machine Learning Research, 6, 1939–1959 http://www.jmlr.org/papers/v6/quinonero-candela05a.html.

Rad, K. R. & Paninski, L. (2010). Efficient, adaptive estimation of two-dimensional firing rate surfaces via Gaussian process methods. Network: Computation in Neural Systems, 21(3-4), 142–168, https://doi.org/10.3109/0954898X.2010.532288.

Rasmussen, C. E. (2003). Gaussian processes in machine learning. In Summer school on machine learning (pp. 63–71).: Springer.

Rowland, D. C., Roudi, Y., Moser, M.-B., & Moser, E. I. (2016). Ten years of grid cells. Annual review of neuroscience, 39, 19–40, https://doi.org/10.1146/annurev-neuro-070815-013824.

Rule, M. & Sanguinetti, G. (2018). Autoregressive point processes as latent state-space models: A momentclosure approach to fluctuations and autocorrelations. Neural computation, 30(10), 2757–2780, https://doi.org/10.1162/neco_a_01121.

Rule, M. E., Schnoerr, D., Hennig, M. H., & Sanguinetti, G. (2019). Neural field models for latent state inference: Application to large-scale neuronal recordings. PLoS computational biology, 15(11), e1007442, https://doi.org/10.1371/journal.pcbi.1007442.

Sargolini, F., Fyhn, M., Hafting, T., McNaughton, B. L., Witter, M. P., Moser, M.-B., & Moser, E. I. (2006). Conjunctive representation of position, direction, and velocity in entorhinal cortex. Science, 312(5774), 758–762, https://doi.org/10.1126/science.1125572.

Savin, C. & Tkacik, G. (2016). Estimating nonlinear neural response functions using gp priors and kronecker methods. Advances in Neural Information Processing Systems, 29 http://proceedings.neurips.cc/paper/2016/hash/8d9fc2308c8f28d2a7d2f6f48801c705-Abstract.html.

Seeger, M. (1999). Bayesian methods for support vector machines and Gaussian processes. Master’s the-sis, Ecole Polytechnique Federale de Lausanne https://infoscience.epfl.ch/record/175479.

Truccolo, W. (2016). From point process observations to collective neural dynamics: Nonlinear Hawkes process GLMs, low-dimensional dynamics and coarse graining. Journal of Physiology-Paris, 110(4), 336–347, https://doi.org/10.1016/j.jphysparis.2017.02.004.

Truccolo, W., Eden, U. T., Fellows, M. R., Donoghue, J. P., & Brown, E. N. (2005). A point process framework for relating neural spiking activity to spiking history, neural ensemble, and extrinsic covariate effects. Journal of neurophysiology, 93(2), 1074–1089, https://doi.org/10.1152/jn.00697.2004.

Virtanen, P., et al. (2020). SciPy 1.0: Fundamental Algorithms for Scientific Computing in Python. Nature Methods, 17, 261–272, https://doi.org/10.1038/s41592-019-0686-2.

Wu, A., Roy, N. A., Keeley, S., & Pillow, J. W. (2017). Gaussian process based nonlinear latent structure discovery in multivariate spike train data. In Proceedings of the 31st International Conference on Neural Information Processing Systems (pp. 3499–3508). http://proceedings.neurips.cc/paper/2017/hash/b3b4d2dbedc99fe843fd3dedb02f086f-Abstract.html.

Yu, B. M., Cunningham, J. P., Santhanam, G., Ryu, S. I., Shenoy, K. V., & Sahani, M. (2009). Gaussianprocess factor analysis for low-dimensional single-trial analysis of neural population activity. Journal of neurophysiology, 102(1), 614–635 http://proceedings.neurips.cc/paper/2008/hash/ad972f10e0800b49d76fed33a21f6698-Abstract.html.

Zhao, Y. & Park, I. M. (2017). Variational latent Gaussian process for recovering single-trial dynamics from population spike trains. Neural computation, 29(5), 1293–1316, https://doi.org/10.1162/NECO_a_00953.

